# In vitro multi-assay cytotoxicity assessment of iron oxide nanoparticles on neural cells – addressing spectrophotometric interference

**DOI:** 10.1101/2025.10.24.684369

**Authors:** I. U. S. Dhillon, R. M. Cavalieri, E. E. Ghadim, S. Huband, J. Liu, P. J. Sadler, J. F. Collingwood, S. E. Bakker, J. Brooks

## Abstract

Iron-rich nanoparticulate matter air pollution exposure is implicated in the pathophysiology of Alzheimer’s disease. The use of colorimetric cell viability assays to ascertain the potential cytotoxicity of iron oxide nanoparticles have highlighted conflicting results in various *in vitro* models, because of spectrophotometric interference. Attempts to resolve this confounding factor to reveal accurate measurements have been insufficient. To address this interference issue, we tested three control methods (readout correction, plate transfer, and above-plate reading) on *in vitro* cytotoxicity measurements via Neutral Red Uptake (NRU), Resazurin, and MTT assays, of iron oxide and magnetite nanoparticulate exposure on SK-N-SH and IMR-32 human neural cell lines. We observed varying degrees of interference based on cell line, nanoparticle type, exposure time, assay type, and the readout method (colorimetric or fluorometric). The readout correction method did not remove interference completely, the plate transfer method tended to cause overestimation of cell viability beyond the original interference, and fluorescence readings from above the plate were less sensitive than the conventional bottom reading. The Resazurin assay with fluorometric readout presented no interference in most conditions investigated in this work, therefore presents advantages over both colorimetric assays and interference control methods. It is recommended to forego colorimetric assays in future studies of nanoparticle cytotoxicity as they are more susceptible to interference.

## 1. Introduction

In 2019, the WHO identified that “99% of the world’s population was living in places where the WHO air quality guidelines were not met”.^1^ This presents an issue for majority of the human population, as air pollution is a ubiquitous environmental risk factor that has epidemiological links with several global health challenges, including lung disease^2^, cancers^3–5^, and most notably neurodegenerative disorders.^6–8^ Recently, Rogowski *et al*. conducted a meta-analysis of the evidence concerning exposure to urban air pollution as a risk factor for the onset of dementia, and found dementia diagnosis to be significantly associated with long-term exposure to PM_2.5_ (airborne particulates with a diameter 2.5 microns or less), nitrogen dioxide, and black carbon.^9^

The UK Heavy Metals Monitoring Network reports iron to contribute up to 1% of PM mass, making it the most abundant metal measured in the environmental sites studied across the UK.^10^ Iron is also a key mineral for normal central nervous system (CNS) development and function.^11^ Disrupted brain iron homeostasis is a well-established feature of Alzheimer’s disease (AD), whilst the amount of iron in the human brain naturally increases with age.^12^ Therefore, the pathophysiological role of iron in neurodegenerative diseases warrants great attention.^13^ Magnetite (Fe(II)Fe(III)_2_O_4_), a redox-active iron oxide, is a significant nanoparticulate pollutant present in urban populations, formed during combustion- and friction-derived reactions such as vehicular wear and tear.^14,15^ Prior research conducted on post-mortem brain tissues from Mexico City residents (a region with high levels of air pollution) found that urban residents with Alzheimer’s disease continuum had higher Fe_3_O_4_ levels in the brain versus clean air controls.^16^ As such, it must be considered whether exposure to iron-rich nanoparticulate matter is a modifiable risk factor for Alzheimer’s disease.

Toxicological assessments of PM exposure on the nervous system are abundant in the literature, particularly regarding iron-oxide nanoparticles (IONPs) *in vitro, in vivo*, and *ex vivo*.^17–22^ Various types have been assessed, including uncoated^23^ and coated (with organic and/or inorganic materials)^24,25^, environmentally derived^19^ and synthetic^26^, as well as mixed- and single-phase samples (e.g., magnetite vs iron-oxide samples).^27^ In the case of in vitro work, cell viability assays based on different biomarkers, such as membrane integrity^28^ or cellular metabolic activity^29,30^, are used to determine the effect of a nanoparticle under a specific set of conditions. They are divided into colorimetric, fluorometric, luminometric, flow cytometric, and dye exclusion assays.^31^ Perhaps the most common colorimetric (the former two) and fluorometric (the latter) assays are Neutral Red Uptake (NRU), 3-[4,5-dimethylthiazol-2-yl]-2,5 diphenyltetrazolium bromide (MTT), and Resazurin.^28,29,32,33^ Prior research using such assays in the context of iron nanoparticulate cytotoxicity has shown conflicting results, where they show significant cytotoxicity in a dose- or time-dependent manner under some conditions, or appear to have increased cell viability instead.^23,34–36^

The increase in cell viability because of nanoparticulate exposure, when this is not the intended result, is reportedly attributable to spectrophotometric interference (SPI), a result of the spectral properties of nanoparticles.^37^ This causes them to have overlapping absorption and emission spectra with the chromophores and fluorophores leveraged in viability assays.^37^ For example, SPI is the reason for the artificial increases in measured viability seen when neuronal and glial cells were exposed to IONPs.^34^ Therefore, interference is a problem that impairs the accuracy and reliability of *in vitro* nanotoxicology assessments, when researching the links between air pollution-relevant nanoparticles and human brain cell models. Several methods to mitigate SPI have been proposed in the literature, however, there is no universal approach.^35,38– 40^ Nevertheless, several methods must be trialled for conventional cytotoxicity tests to ascertain which one enables data reliability. Researching SPI control methods would not just benefit nanotoxicology workflows alone. Ascertaining the non-cytotoxic concentrations of IONPs can drive further research into uptake, internalisation, and intracellular dynamics for example, without affecting the viability of the biological model.

The present study aims to employ the conventional *in vitro* cytotoxicity tests (NRU, Resazurin, MTT) to assess the effects of two iron-rich nanoparticle systems – mixed-phase iron oxide nanoparticles and magnetite nanoparticles, on the human neuroblastoma cell lines SK-N-SH (epithelial) and IMR-32 (fibroblast). A multi-assay combinatorial approach was used to compare results from different routine assays. The efficacy of three interference control methods – interference subtraction^34^, plate transfer^41^, and above plate fluorescence reading, is evaluated in parallel. Furthermore, the behaviour of the particles with all the assay components is assessed to understand when interference occurs. This study will therefore evaluate whether any of the tested methods can accurately determine the real cytotoxicity of mixed-diameter iron-oxide nanoparticles. Investigating the interactions between iron-rich nanoparticulate matter, such as magnetite, and human brain cells may enable better understanding of the mechanisms which underpin toxicological links between PM and neurodegeneration. This will inform the development of environmental policies to help mitigate exposure to air pollution and the prevalence of neurodegenerative pathologies.

## 2. Experimental

### Materials

SK-N-SH and IMR-32 cells were from the American Type Culture Collection (ATCC). Eagle Minimum Essential Medium (EMEM), both with and without phenol red, dimethyl sulfoxide (DMSO), glacial acid 100%, ethanol absolute, Triton X-100 (for molecular biology), Trypan Blue solution, Neutral Red Uptake (NRU) assay reagent, and Resazurin *in vitro* toxicology assay kit were all from Sigma Aldrich. Foetal bovine serum (FBS) was from Labtech. Trypsin ethylenediaminetetraacetic acid (EDTA; 0.25%/1 mM) solution, penicillin/streptomycin solution, and sterile HEPES (0.1 M) buffer solution were prepared in-house and used after thawing if necessary. Dulbecco’s Phosphate Buffered Saline (PBS) was from Cytiva Life Sciences. CyQUANT™ MTT cell viability assay kit was from Invitrogen (Thermo Fisher Scientific). 96-well optical-bottom black and transparent cell culture plates (with low evaporation lids) were from Fisher Scientific. 75 cm^2^ cell culture flasks were from Corning Incorporated.

### Cell Culture

The SK-N-SH neuroblastoma cell line is derived from a tumour of a female patient and has epithelial morphology.^42^ The IMR-32 neuroblastoma cell line is derived from a tumour of a 13-month-old male patient and consists of two cell types - small neuroblast-like cells and large hyaline fibroblast cells.^43^ Both cell lines are adherent. SK-N-SH and IMR-32 cells were cultured in EMEM cell culture medium containing 10% FBS, 1% penicillin G (100 U/mL), 1% streptomycin sulphate (100 μg/mL), 1 g/L glucose, and 2mM L-glutamine. Cells were maintained at 37 ºC in a humidified atmosphere of 5% CO_2_. To harvest cells for experimentation or cell splitting, trypsin/EDTA solution was used for detachment, and a Neubauer counting chamber was used for cell counting.

### Nanoparticle preparation

The uncoated iron oxide nanoparticles (IONPs) were prepared in-house according to a confidential synthetic protocol. The stock powder was suspended in 99.5% ethanol, and the concentration was determined by thermogravimetric analysis. Before each experiment, a 300 μg/ml suspension was prepared in 0.1 M HEPES aqueous buffer (to reduce IONP aggregation) and sonicated in a water bath for 10 min. Serial dilutions were carried out to obtain the different treatment concentrations.

Magnetite nanoparticles (Fe_3_O_4_, MNPs) were synthesised based on previous reported co-precipitation methods.^44,45^ Briefly, Milli-Q water (10 mL) was degassed with N_2_ for 30 min at 27 °C followed by the addition, under inert atmosphere, of FeCl_2_ (0.32 g, 2.5 mmol) and FeCl_3_ (0.83 g, 5 mmol) to give a final ratio of Fe(II)/Fe(III) = 0.5. Under continuous stirring, ammonia solution (65 mL, 1.5 M) was slowly added (50 μL/sec) into the reaction mixture, and a black solid began to form in solution. After the base addition was complete, the black reaction mixture was left stirring for 1 h. The MNPs were separated by attracting them towards a magnet and washed with water (10 x 100 mL), isopropanol (2x 30 mL), and acetone (2x 30 mL). Finally, they were dried under vacuum for 6 h (92% yield). The concentration was determined by thermogravimetric analysis. Before each experiment, a 1100 μg/ml suspension was prepared in complete EMEM, vortexed, sonicated for 5 min, then vortexed again. Serial dilutions were carried out to obtain the different treatment concentrations.

### Nanoparticle characterisation

Transmission electron microscopy (TEM) was used to determine the primary particle size and morphology using a JEOL2100Plus microscope equipped with a Gatan OneView IS camera or a JEOL ARM200F microscope equipped with a Gatan Orius SC1000 CCD camera, both operated at 200 kV. Images were analysed using ImageJ. Powder X-Ray Diffraction (PXRD) was used to confirm which iron oxide phases were present in the sample post-synthesis. PXRD experiments were conducted on Anton Paar XRDynamic 500 equipped with a Primux 3000 X-ray tube giving Co K*α1,2* radiation (*λ* = 1.7902 Å) and a Pixos 2000 1D detector. The sample was grounded and mounted in the Anton Paar TTK 600 stage. Each scan was recorded with samples spinning, at a step size of 0.01° and within an angular range of 10.0–95.0° (2*θ*). Dynamic light scattering (DLS) was used to determine the average hydrodynamic diameter and zeta potential of the nanoparticles using an Anton Paar Litesizer 500 with a 40 mW 658 nm laser, at an automatic or backscatter (175º) angle. Each sample was sonicated for a further 1 min before taking the DLS measurements. Ultraviolet-visible (UV-Vis) spectroscopy was used to determine the absorbance as a function of concentration of the nanoparticles using an Agilent Cary 60 UV-Vis spectrophotometer.

### Exposure conditions

For each assay experiment, 7 different concentrations (100, 50, 25, 12.5, 6.25, 3.13, and 1.56 μg/ml), a negative, positive, and vehicle control (if dispersed in 0.1 M HEPES), and up to two exposure periods (4 and 24 h) were evaluated. Cell culture medium without nanoparticles was used as the negative control. Triton X-100 (1% v/v) was used as the positive control. For the vehicle control, the same volume of 0.1 M HEPES without IONPs was added to the control wells.

### Neutral red uptake (NRU) assay

The NRU assay was carried out as previously described by Repetto *et al*.^28^ SK-N-SH and IMR-32 cells were seeded in separate 96-well plates at 25,000 cells/well and 30,000 cells/well in 100 μL of complete medium and then incubated for 24 h at 37 ºC/5% CO_2_, respectively. After 24 h, the medium in the wells was replaced and the cells were exposed to the IONPs for 24 h or the MNPs for 4 or 24 h. After incubation, the medium was removed, and the cells were washed with PBS (150 μL/well). NR reagent (40 μg/ml) was prepared in phenol-red free complete EMEM and was added (100 μL/well) to the cells followed by incubation for 3 h. After that, the cells were washed with PBS (150 μL/well), and then NRU destain solution (50% ethanol absolute, 49% deionised water, 1% glacial acetic acid; 150 μL/well) was added to elute the dye from the cells. The plate was shaken for 15 min to form a homogenous solution, then left to sit for 5 min. Then absorbance was measured at 540 nm and fluorescence was measured at an excitation and emission wavelength of 530 nm and 645 nm respectively, using a Tecan Infinite M Nano^+^ microplate reader. To correct for the spectrophotometric interference, the absorbance values of the IONPs in destain solution were used. For the MNPs, 50 μL from each well was transferred to a new plate and the absorbance was re-measured.

### Resazurin assay

The resazurin assay was carried out as previously described by Larsson and Parris.^46^ SK-N-SH and IMR-32 cells were seeded in separate 96-well plates at 30,000 cells/well and 40,000 cells/well in 100 μL of complete medium respectively and then incubated for 24 h at 37 ºC/5% CO_2_. After 24 h, the medium was replaced, and the cells were exposed to the IONPs for 24 h or the MNPs for 4 or 24 h. After incubation, the medium was removed, and the cells were washed with PBS (150 μL/well). A 10% resazurin solution was prepared in phenol red-free complete EMEM and was added to the cells (100 μL/well). The SK-N-SH and IMR-32 cells were incubated for 2 and 3 h respectively; the plates were wrapped in foil. After incubation, absorbance was measured at 570 nm with a reference wavelength of 630 nm, and fluorescence was measured with excitation and emission wavelengths of 560 and 590 nm respectively, using a Tecan Infinite M Nano^+^ microplate reader. To correct for the spectrophotometric interference, the absorbance and fluorescence values of the IONPs in 10% resazurin solution were used. For the MNPs, fluorescence measurements were taken from above and below the plate and compared.

### MTT assay

The MTT assay was carried out as described in the assay kit protocol. SK-N-SH and IMR-32 cells were seeded 96-well plates at 25,000 cells/well and 45,000 cells/well in 100 μL of complete medium and then incubated for 24 or 48 h at 37 ºC/5% CO_2_ respectively. After 24 - 48 h, the medium was replaced, and the cells were exposed to the MNPs for 4 or 24 h. After incubation, the medium was removed, the cells were washed with PBS (150 μL/well), then phenol-red- and serum-free EMEM was added to the wells (100 μL), along with MTT solution (5 mg/ml prepared in sterile PBS; 10 μL). The SK-N-SH and IMR-32 cells were incubated for 3 and 2 h respectively; the plates were wrapped in foil. After incubation, the medium was removed (85 μL) carefully so as not to disturb the formazan crystals, and DMSO (100 μL) was added and pipetted thoroughly in the wells to solubilise the formazan. Then the plate was shaken at 37 ºC for 10 min before the absorbance was measured at 540 nm with a reference wavelength of 630 nm.

### Nanoparticle spectrophotometric interference studies

To assess possible interference that the nanoparticles may have had on the cytotoxicity assay procedures, cell-free experiments were conducted (1) to measure the optical properties of nanoparticles in the cell media, and (2) to measure the effect of the presence of nanoparticles with the assay components.

#### (1) Measurement of optical properties in media

A 96-well transparent cell culture plate was loaded with 100 μL medium/well. A serial dilution of IONPs or MNPs were prepared in their respective media (0.1 M HEPES or complete EMEM) and added to the plate of medium (50 or 10 μL treatment/well, respectively). This was incubated at 37 ºC/5% CO_2._ The absorbance at 540 nm was measured at 4 and 24 h to reflect the treatment incubation times tested.

#### (2) Effect of presence of nanoparticles with assay components

The NRU, Resazurin, and MTT assay reagents were incubated with the nanoparticles in the absence of cells. For the NRU and MTT assays, the nanoparticles were incubated with the respective NRU and MTT reagents (dissolved in the same medium as used in the cell assay experiments) for 3 h at 37 ºC/5% CO_2_ and the absorbance was measured at 540 nm. The nanoparticles were also dispersed in the dissolving agents for the assays (destain solution and DMSO respectively), and the absorbance at 540 nm was measured immediately. For the Resazurin assay, the nanoparticles were incubated in a 10% resazurin solution (prepared as above), and the fluorescence was measured at 2 and 3 h.

### Statistical analysis

All statistical analyses were performed using GraphPad Software LLC (version 10.5.0). The cytotoxicity assay data were expressed as the mean ± SEM of three independent experiments, each carried out in three (MTT) or six replicates (NRU, Resazurin) per condition. Statistical analyses were performed by one-way ANOVA followed by Dunnett’s test, or by Kruskal-Wallis test followed by Dunnett’s test for non-normal data. Only *p* values less than 0.05 were considered as significant.

## 3. Results & Discussion

### 3.1. Nanoparticle characterisation

Table 1 shows the main physicochemical characteristics of the IONPs and MNPs investigated in this work – primary particle size and morphology, hydrodynamic diameter, and zeta potential The DLS measurements were taken in 0.1 M HEPES buffer, deionised (DI) water, phenol red-free complete and incomplete cell medium (EMEM). TEM image analysis evidenced that the IONPs (Figure **1a**) were angular with sharp edges and had an average diameter of 8.06 nm (see figure S**6a-c** for the TEM images used to ascertain diameter of the spherical nanoparticles). Furthermore, the MNPs had a spherical morphology, with a mean diameter of approximately 15 and 18 nm according to TEM and SAXS, respectively.

**Figure 1.**
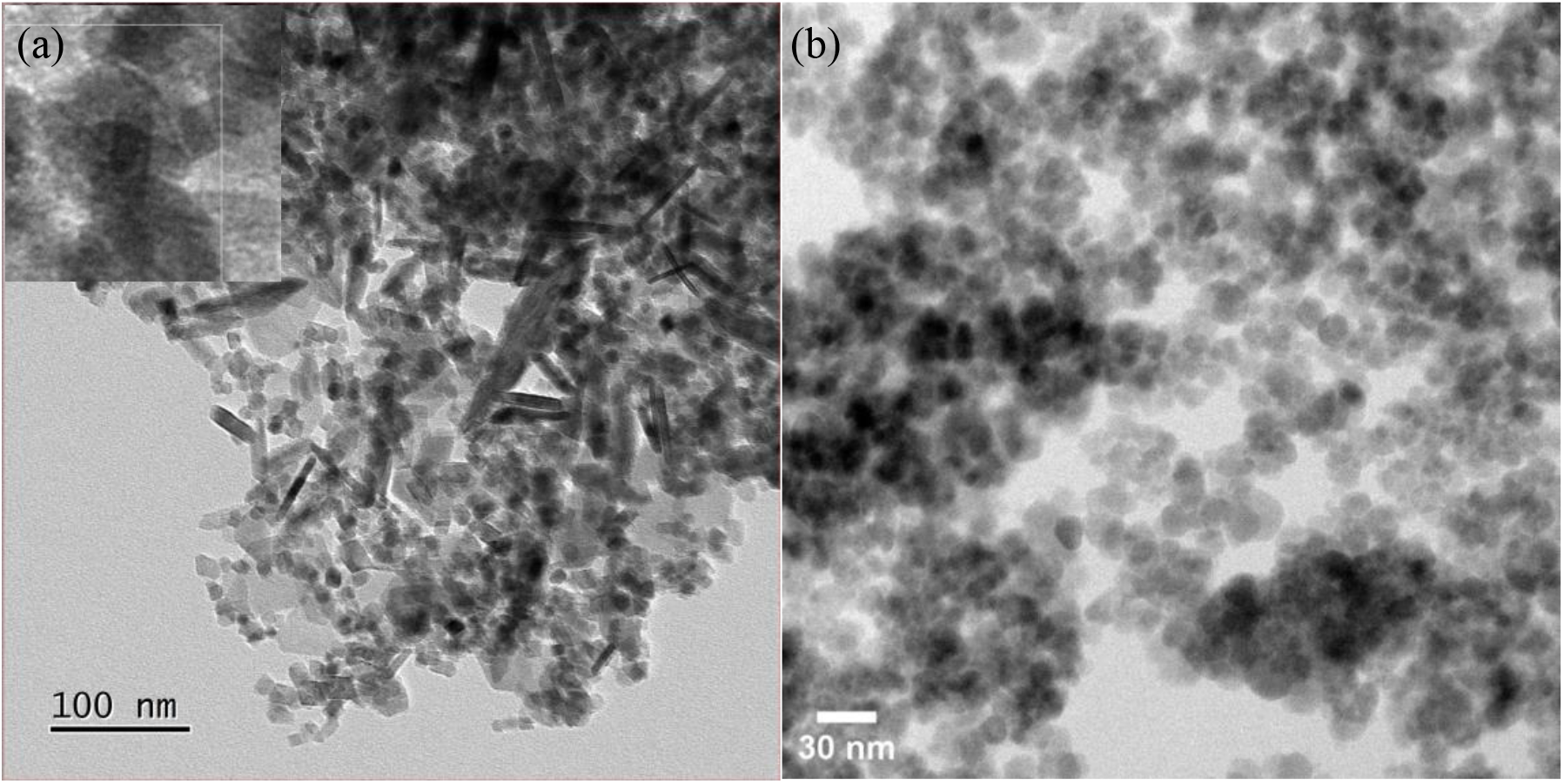
Transmission electron microscopy images of the iron-rich nanoparticles tested in this study. (a) Inset image of the IONPs at a 1 μg/ml concentration in EtOH at 30 nm scale; (b) image of the MNPs at a 10 μg/ml concentration in EtOH at a 30 nm scale.

**Table 1.**
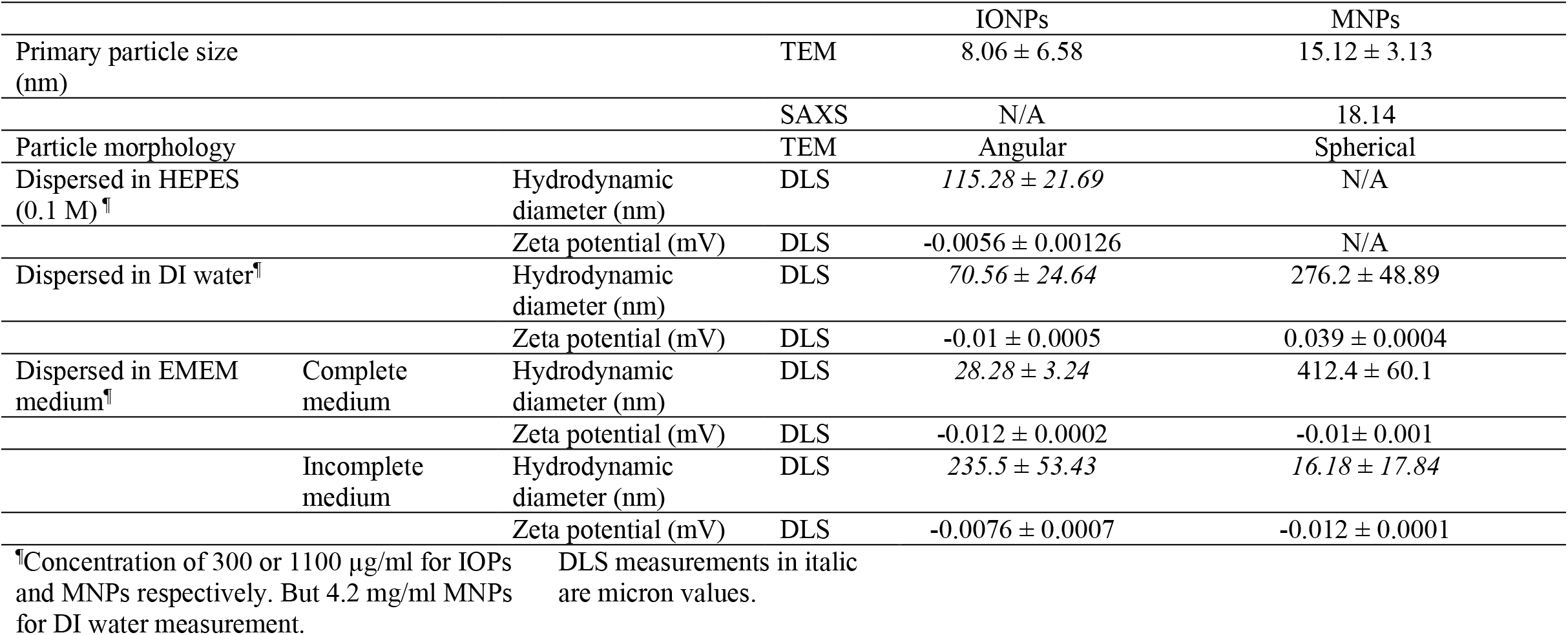
Physicochemical characteristics of the iron oxide particles and magnetite nanoparticles.

The average hydrodynamic diameters (HD) of the IONPs in four different media – deionised (DI) water, 0.1 M HEPES, phenol red-free complete and incomplete medium, were all in the micron size range, implying large coronas formed around the IONPs in solution. At 300 μg/ml the HD was larger in HEPES (115.28 μm) compared to DI water (70.56 μm), and in incomplete medium (235.5μm) compared to complete medium (28.28 μm). The HD values of the MNPs in DI water and complete medium were approximately 276 and 412 nm respectively, whereas in incomplete medium it was 16.18 μm. HEPES buffer serves as a stabiliser to prevent the aggregation of metal nanoparticles as the molecules attach to the surfaces of the particle cores, encouraging electrostatic repulsion between the particles.^47^ Therefore, since HEPES molecules are larger than water molecules, this leads to a larger size being measured in the solution. In addition, complete and incomplete medium differ in their composition, affecting the HD too. For example, incomplete medium lacks serum proteins and may have higher free ion availability because of this, whereas complete medium has been supplemented with foetal bovine serum. As such, the presence of serum proteins can change the intermolecular forces that nanoparticles are subjected to by forming a biomolecular corona, which influences their in vitro behaviour (e.g., nanoparticle uptake, propensity for agglomeration, and potential cytotoxicity).^48^

This mostly corroborates with the average zeta potential measurements, where in the complete medium, the magnitude is greater (more negative) for the IONPs as the serum proteins adsorb to the surface, and it is likely are presenting negatively charged groups outwards from the core. As such, there would be a more negative net surface charge, decreasing aggregation (hence the smaller HD), leading to the greater colloidal stability in complete medium due to the protein coronas forming. Despite any marginal differences in colloidal stability, the zeta measurements were much lower than the ± 30 mV “highly stable” threshold, suggesting there is much aggregation behaviour of these IONPs, which fits with the micron hydrodynamic diameter measurements obtained.^49^ As with the IONPs, the zeta potentials of the MNPs were much lower than the ± 30 mV threshold, suggesting a lack of colloidal stability, but not to the same severity as the IONPs, since the HD values measured were smaller in each medium comparatively.

### 3.2. Optical properties of the particles in solution

The optical absorption spectra of the IONPs and MNPs were determined by UV-Vis spectroscopy, and their behaviour in the presence of complete EMEM medium following the same incubation time points and at the same wavelengths as used in the in vitro cytotoxicity assays were assessed as well (Figure **2**). The UV-Vis absorbance was measured between 800 and 250 nm of the IONPs and MNPs, dispersed in ethanol and water at 100 μg/ml respectively. At 800 nm, the IONPs had high absorbance, starting at 0.97, which increased consistently as the wavelength decreased, reaching a plateau at approximately 380 nm, where the absorbance was approximately 2.80. In contrast, the absorbance of the MNPs began close to 0 at 800 nm and increased at a smaller gradient than the IONPs. The absorbance only became greater than 1 at a wavelength of approximately 320 nm, and did increase, albeit slowly, up until 250 nm. According to the Beer-Lambert Law, absorbance is proportional to the molar absorption coefficient, path length, and concentration. Since the latter two are identical in both measurements, this suggests the stark difference in absorbance seen depends on the different molar absorption coefficients of magnetite in solution and a mixed-phase iron oxide solution, where the coefficient is based on chemical composition. The high absorbance across a wide range of wavelengths implies that the particle samples may present spectrophotometric interference in the in vitro cytotoxicity assays, particularly at 540 nm which is the measurement wavelength for the MTT and NRU assays.

**Figure 2.**
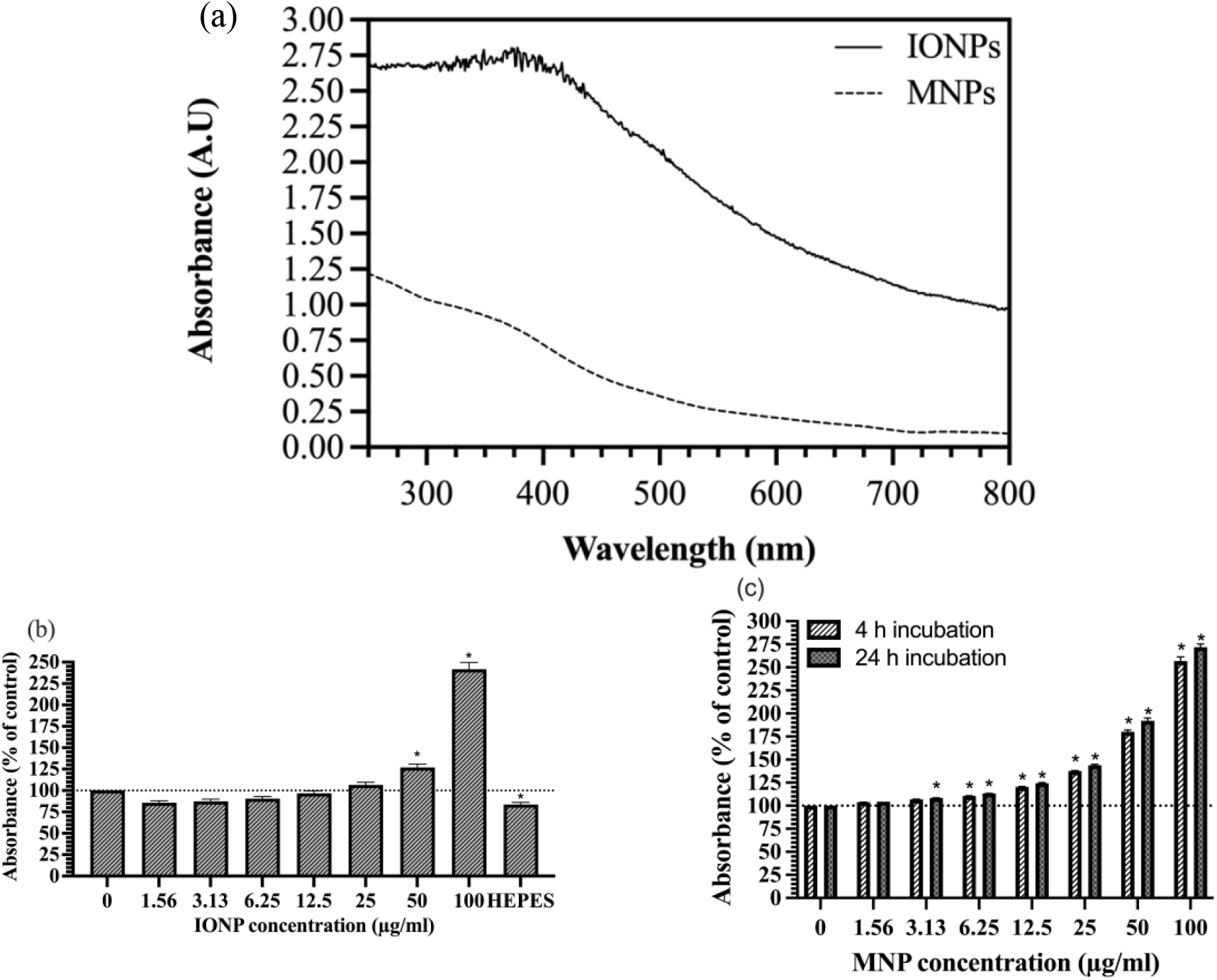
(a) UV-Vis spectra of the particles tested in this study. The black solid line represents the IONPs at 100 μg/ml dispersed in ethanol absolute and the black dashed line represents the MNPs at 100 μg/ml dispersed in deionised water. (b,c) Absorbance at 540 nm (as used in the NRU and MTT assays) of the IONPs and MNPs in complete EMEM medium following a 4- or 24-hour incubation.

Before conducting the cytotoxicity tests, the NPs were incubated with the complete EMEM medium to investigate whether their optical properties as seen in UV-Vis would present an interference effect on the absorbance readings. The IONPs were dispersed in HEPES buffer to reduce aggregation and were then added to wells containing complete medium, whilst the MNPs were dispersed in the medium and then added to wells containing the same medium. Due to the difference in experimental procedure, we see differing results. With the IONPs, the results showed that only at 50 and 100 μg/ml concentration the absorbance reading was significantly greater than the control (no IOPs), while at 25 μg/ml it was marginally but not significantly greater than the control. Below 25 μg/ml the absorbance appeared to be lower than the control, which is contradictory. However, it is because when the IONPs were dispersed in HEPES in a serial dilution, the lower concentrations of IONPs do not appear as a dark solution (HEPES is a colourless buffer). Therefore, adding an essentially colourless solution (with a low concentration of IONPs) to wells containing phenol red-containing medium diluted the colour, resulting in lower absorption compared to the control. On the other hand, when the MNPs were incubated with the medium and the absorbance measured after two time points, the results showed that starting at 3.13 μg/ml the absorbance readings were significantly greater than the respective control at 24 h, and then for each consecutive concentration at both time points as well. At 100 μg/ml, the absorbance increased 2.56-fold and 2.71-fold at 4 and 24 h respectively. As such, it was expected that the NP samples would display spectrophotometric interference in the in vitro cytotoxicity assays if they remained in the wells following washing steps in the experimental procedure. Furthermore, the NPs were also incubated with the assay reagents MTT, NRU, and Resazurin, as well as the dissolving agents DMSO and destain solution (50% ethanol absolute, 49% deionised water, and 1% acetic acid), similar results were seen with MTT reagent and the dissolving agents (see SI figure **9**). Previous literature has identified spectrophotometric interference by IONPs coated with inorganic (silica) and organic materials (e.g., oleic acid) in the NRU and MTT assays (540 nm).^34,35^ As such, it may be difficult to accurately quantify the potential cytotoxicity of iron-rich nano- and micron-sized particulate matter without appropriate methods to control for interference. A possible method to mitigate interference is to functionalise the surfaces of NPs, to increase steric and/or electrostatic repulsion. However, this would change the physicochemical characteristics, and therefore the NP system would deviate further from what it’s trying to represent.^48^

Instead of changing the nanoparticle’s surface structure to mitigate against interference, this work instead applied several interference control methods as an alternative means to unmask the cytotoxicity of iron-rich NPs. These were: (1) performing cell-free IONP or MNP assays in the relevant media (destain solution for NRU or 10% Resazurin solution) and subtracting the absorbance of each concentration from the cell viability data to assess whether the interference is eradicated; (2) conducting plate transfer whereby the same volume from each well of the assay plate is transferred to a clean plate without touching the bottom of the wells and the measurements are re-run; (3) reading fluorescent assays from above and below the plate and comparing the readouts.

### 3.3. In vitro cytotoxicity of the IONPs and MNPs

After characterising the nanoparticles and optimising the parameters for the in vitro cytotoxicity assays, three classical tests were performed – NRU, Resazurin, and MTT. The SK-N-SH and IMR-32 cells were exposed to IONPs for 24 hours, and then viability was assessed using the first two tests. The key results are presented in Figure **4**, both the original and corrected data sets are shown as an evaluation of the first interference control method.

**Figure 4.**
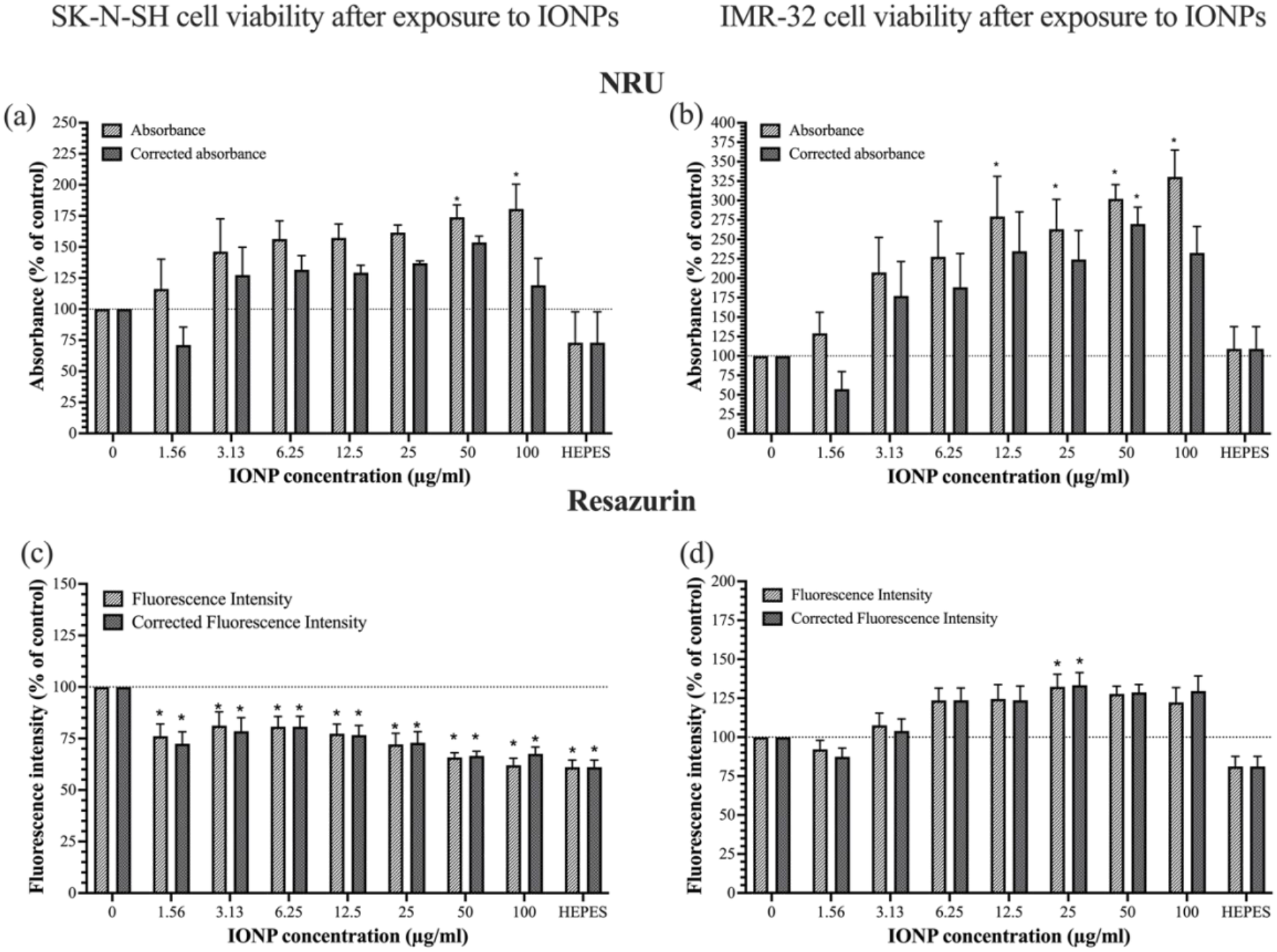
Main cell viability diagrams of the SK-N-SH and IMR-32 cell line exposures to IONPs for 24 hours. (a,b) Viability assessment of SK-N-SH and IMR-32 cells by NRU with absorbance correction; (c,d) viability assessment of SK-N-SH and IMR-32 cells by Resazurin with fluorescence correction. Bars represent standard error of the mean. Values were normalised to the positive and negative control, considering them as 0 and 100% respectively. \**p* < 0.05 represents a significant difference with respect to the negative control. The remaining assay data, showing the fluorescence measurements and absorbance measurements of the NRU and Resazurin assays respectively are in the supporting information.

For both cell types, after 24 h exposure, the NRU assay showed an apparent concentration-dependent increase in viability when measured by the optical density method, ranging from 116% to 180% at 4.7 - 300 μg/ml for SK-N-SH cells, and from 129% to 330% at 4.7 – 300 μg/ml for IMR-32 cells. When measured by the fluorescent method, the apparent increase in cell viability is less enhanced; for SK-N-SH cells it ranged from 112% to 135% at 4.7 - 18.8 μg/ml, then down to 124% at 300 μg/ml. For IMR-32 cells, it went from 122% to 213% at 4.7 – 37.5 μg/ml, then down to 191% at 300 μg/ml. In both cases, more artificial increase in absorbance was observed for the IMR-32 cells.

Regarding the resazurin assay, similar interference was observed in both the optical density and fluorescence methods for IMR-32 cells, the absorbance generally increased as a function of IONP concentration compared the control. But for SK-N-SH cells, there was no interference. Only when measuring fluorescence did the concentrations of IONPs cause a significant decrease in viability compared to the control (as measuring fluorescence is noted to be more sensitive than absorbance). The viability decreased to 76% and 62% at 4.7 and 300 μg/ml respectively. However, the 0.1 M HEPES control also elicited a significant decrease in viability, perhaps due to the higher than usual molar concentration of HEPES, which may have negatively impacted the cells. Because of this, the remaining data in Figure **4b** cannot be evaluated in the context of whether the IONPs did cause significant cytotoxicity to SK-N-SH epithelial cells, despite significant decreases having been observed. Thus, it was decided that although HEPES can reduce NP aggregation, when testing the MNPs, complete EMEM medium would be used as the dispersant instead from hereon.

However, comparing the NRU to Resazurin data for SK-N-SH cells, there was no interference seen in the Resazurin assay, since the IONPs are not autofluorescent, suggesting that fluorescence-based assays present advantages over absorbance-based assays. However, interference was observed for IMR-32 cells. Looking at the cell-free IONP assays incubated in

Resazurin to determine whether resazurin is reduced at the redox-active surfaces of IONPs to resofurin (thus appearing as “increased viability”), there were no drastic increases in fluorescence (see SI figure **9b**). Therefore, the reason for interference in the Resazurin assay with IMR-32 cells is still undetermined.

To correct for the artificial increases in absorbance due to interference, the absorbance of the IONPs at the same concentration range incubated in destain solution (for the NRU assay) and 10% Resazurin solution were calculated and then subtracted from the original data to determine the “real” absorbances post-exposure. In the NRU assay for both cell types and Resazurin assay for IMR-32 cells, only at 4.7 μg/ml of IONPs did the absorbance correction reduce the viability to below the control. However, it was not significantly different according to 1-way ANOVA. In the Resazurin assay, for SK-N-SH cells, despite already having shown significant cytotoxicity, the method was still trialled. Unexpectedly, at 25 – 100 μg/ml of IONPs, fluorescence subtraction caused the viability to artificially increase, due to the fluorescence measured in the cell-free IONP-Resazurin assay. Since the method was unable to eradicate the interference well, it was assumed to be an unreliable method.

The same characterisation and interference assessment procedure was conducted before testing the potential in vitro cytotoxicity of the MNPs in the same cell lines using all three classic tests – NRU, MTT, and Resazurin, following a 4- and 24-hour exposure. Figure **5** highlights the key results (24-hour exposure) and the remainder are in the Supporting Information. The other two interference methods were assessed in this set of experiments. Regarding the NRU assays, for both cell lines, artificial increases in viability (due to interference) were seen for every concentration of MNP except at 1.56 μg/ml. Although, the interference was less pronounced compared to that obtained in the IOP assays. For example, in the NRU assay for IOPs and SK-N-SH cells, the highest absorbance obtained was 180%, whereas with the MNPs and SK-N-SH cells it was 109%, much closer to the negative control of 100%. On the other hand, with IOPs and IMR-32 cells, the highest absorbance was 330%, but with the MNPs, it was 150%, a 2.2-fold decrease at the same treatment concentration.

**Figure 5.**
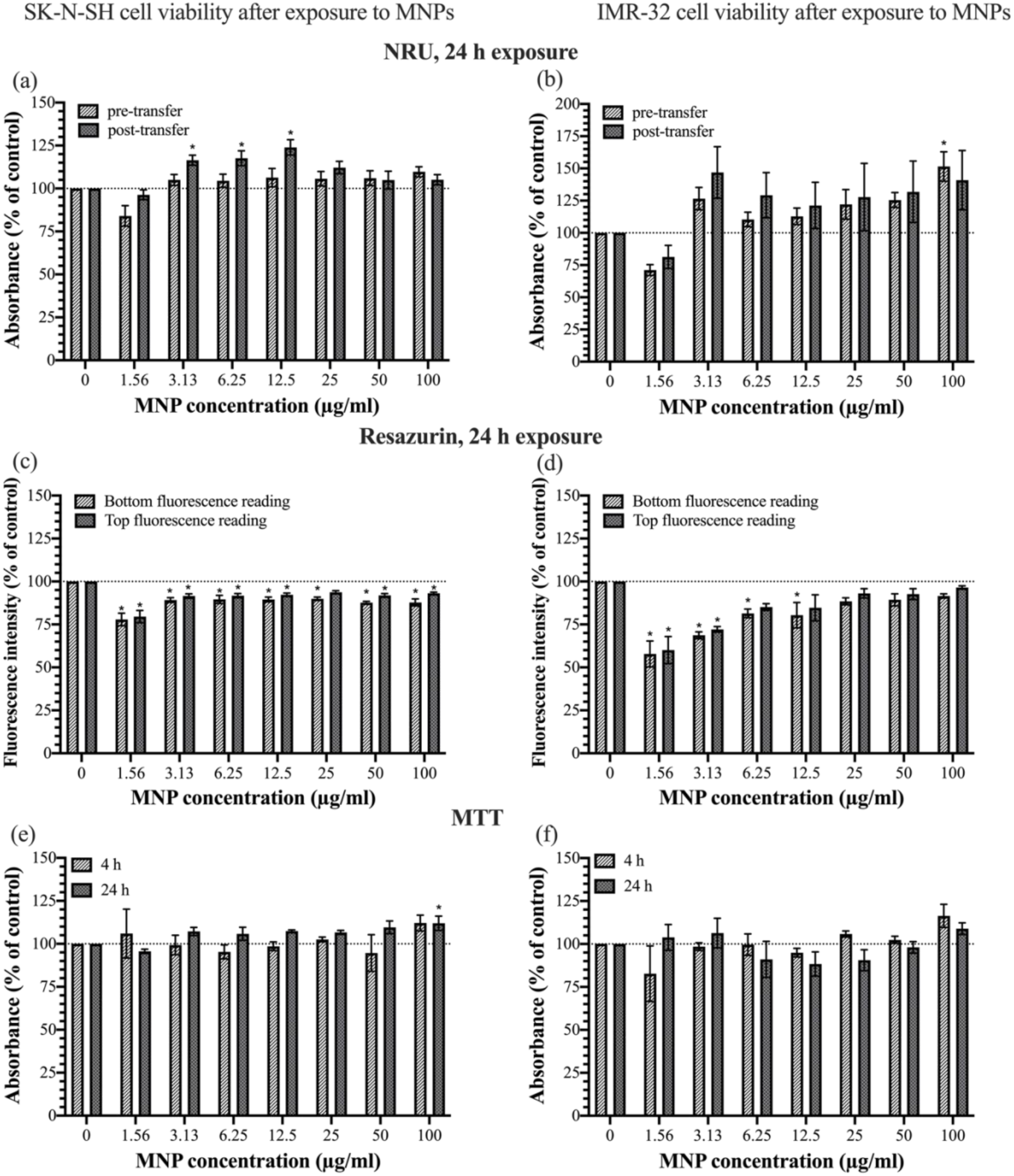
Main cell viability diagrams of the SK-N-SH and IMR-32 cell line exposures to MNPs for 4 and 24 hours. (a,b) Viability assessment of SK-N-SH and IMR-32 cells by NRU with pre- and post-transfer absorbance measurements; (c,d) viability assessment of SK-N-SH and IMR-32 cells by Resazurin with both reading modes; (e,f) viability assessment of SK-N-SH and IMR-32 cells by MTT. Bars represent standard error of the mean. Values were normalised to the positive and negative control, considering them as 0 and 100% respectively. \**p* < 0.05 represents a significant difference with respect to the negative control. The remaining assay data, showing the NRU and Resazurin assays following 4-hour incubation with MNPs, are in the supporting information.

Longhin *et al*. proposed the plate-transfer method as a control method in the context of assessing the potential cytotoxicity of several non-iron containing nanomaterials using the Resazurin assay, so its reproducibility was assessed here.^41^ A consistent volume of NRU solution from each well of the original plate was added to a new plate to which no MNPs were added (and could have stuck to the well bottom surface). The new plates were also read at the same wavelength (540 nm). At every concentration except for 100 μg/ml (and 50 μg/ml with the SK-N-SH cells), plate transfer also led to artificial increases in absorbance (after normalisation to both controls), which was unexpected. For example, at a 12.5 μg/ml treatment with SK-N-SH cells, the absorbance increased from 106% to 123%. It was thus assumed that this method was not suitable for the specific set of conditions used here – SK-N-SH and IMR-32 cells exposed to < 20 nm uncoated synthetic MNPs.

## Conclusion

In conclusion, this research provides reinforcing insights regarding the challenge of addressing spectrophotometric interference during in vitro toxicology assessments of iron-oxide nanoparticles. The key insights regarding the control methods investigated are as follows. First, the readout correction method using data from cell-free NP-containing assays did not completely remove interference in every instance, so lacks universal efficacy. Second, the plate transfer method caused overestimation of cell viability values beyond the initial data, so also lacks efficacy. Third, the above-plate fluorometric reading is less sensitive than the conventional below-plate reading, therefore unnecessary to perform. Lastly, depending on cell type and exposure time, the Resazurin fluorometric assay is not confounded by interference, so can accurately determine the cytotoxicity of the nanoparticle of interest.

While this study focussed on iron-oxide nanoparticulate matter exposure on human neural cells, the implications go beyond this specific system, to other metal-containing nanoparticles and various mammalian cell types. Spectrophotometric interference is not unique to iron, and as such, the main insight that fluorometric assays such as Resazurin evade this issue, whereas colorimetric assays may not, should be extended to other investigations. The question now lies in which other fluorometric assay workflows can be investigated to ascertain a multi-fluorometric-assay approach to investigating nanoparticulate cytotoxicity. Further investigation into other robust, reproducible, and cost-effective assays is required.

## Acknowledgements

The authors would like to acknowledge the University of Warwick Research Technology Platforms (Advanced Bioimaging Polymer Characterisation, and X-Ray Diffraction) for assistance in the research described in this paper. The authors would also like to acknowledge Nina Pučeková, PhD candidate of the McAinsh Group at the University of Warwick, for assistance with the nanoparticle size analysis of the TEM images of the nanoparticles. I.U.S.D was supported by funding through the UKRI, J.B was supported by the Race Against Dementia fellowship.

## Conflicts of interest

The authors declare no conflicts of interest.

## Notes

### Competing Interest Statement

The authors have declared no competing interest.

