## Supplementary Information for "In vitro multi-assay cytotoxicity assessment of iron oxide nanoparticles on neural cells – addressing spectrophotometric interference"

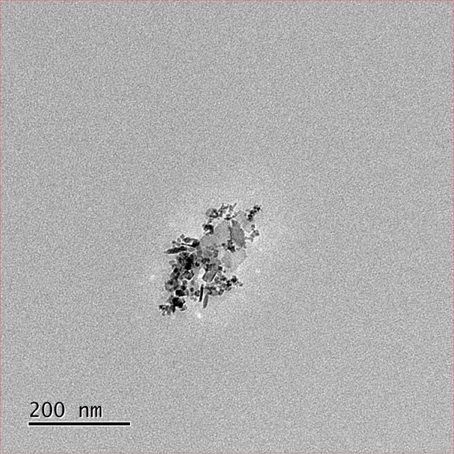

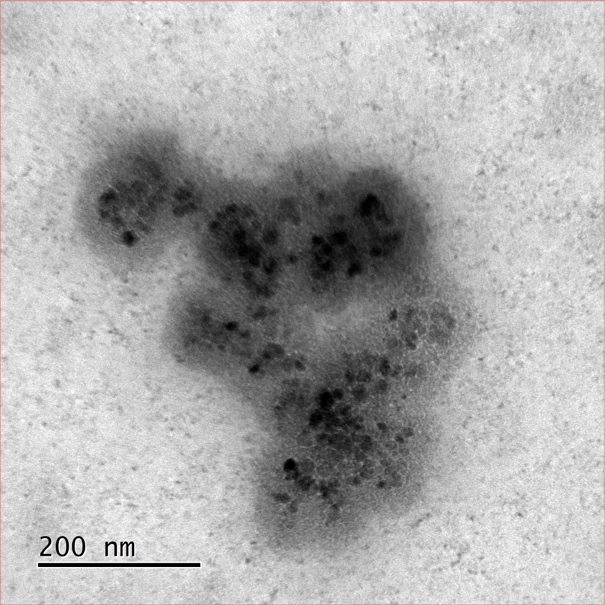

**Figure 6**. Transmission electron microscopy images of the iron-rich nanoparticles tested in this study. (a-c) The images of IONPs dispersed in ethanol used to measure the primary particle size. (d) Image of the IONPs at a 1 µg/ml concentration dispersed in 0.1 M HEPES buffer; (b) image of the MNPs at a 10 µg/ml concentration dispersed in complete EMEM medium.

(d)

(e)

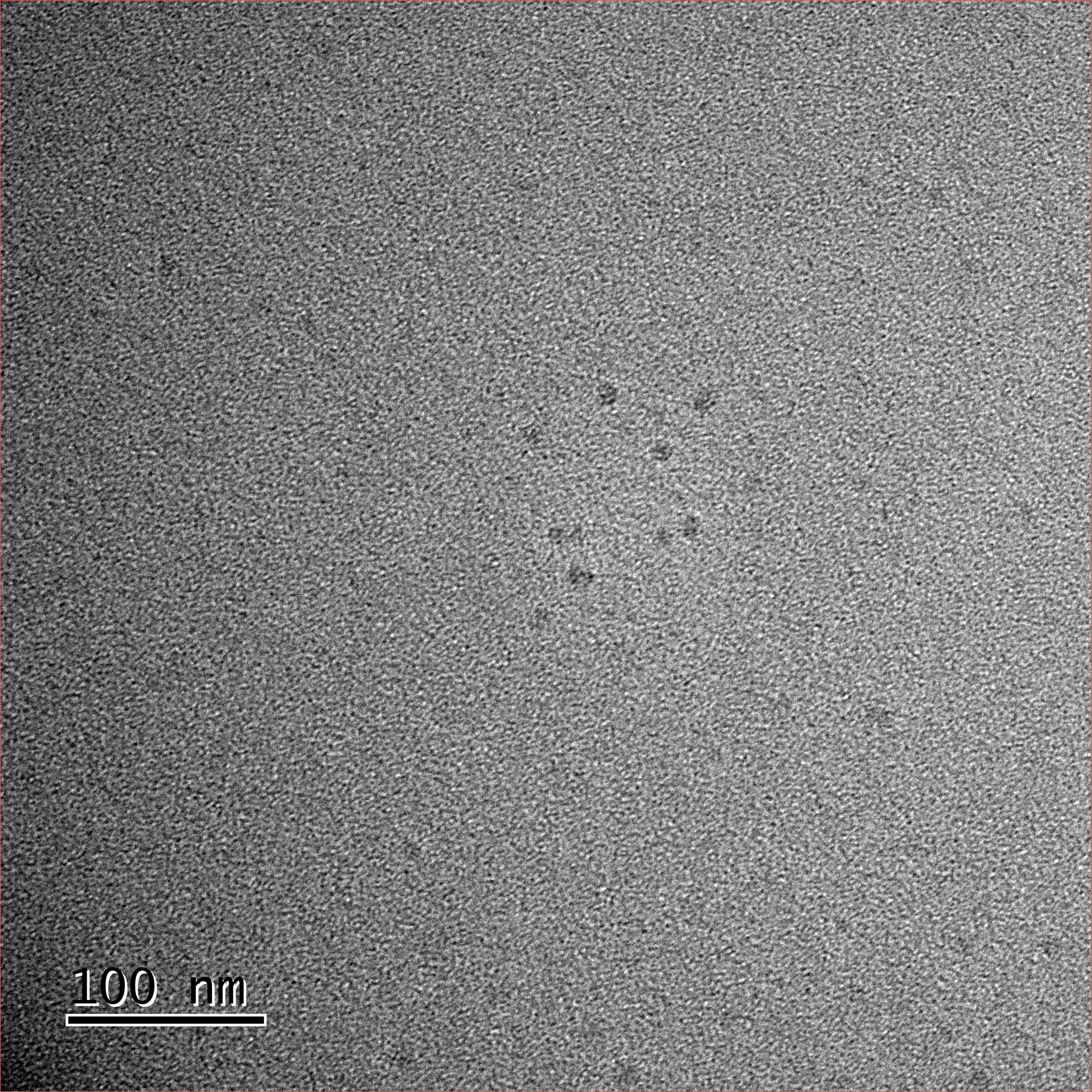

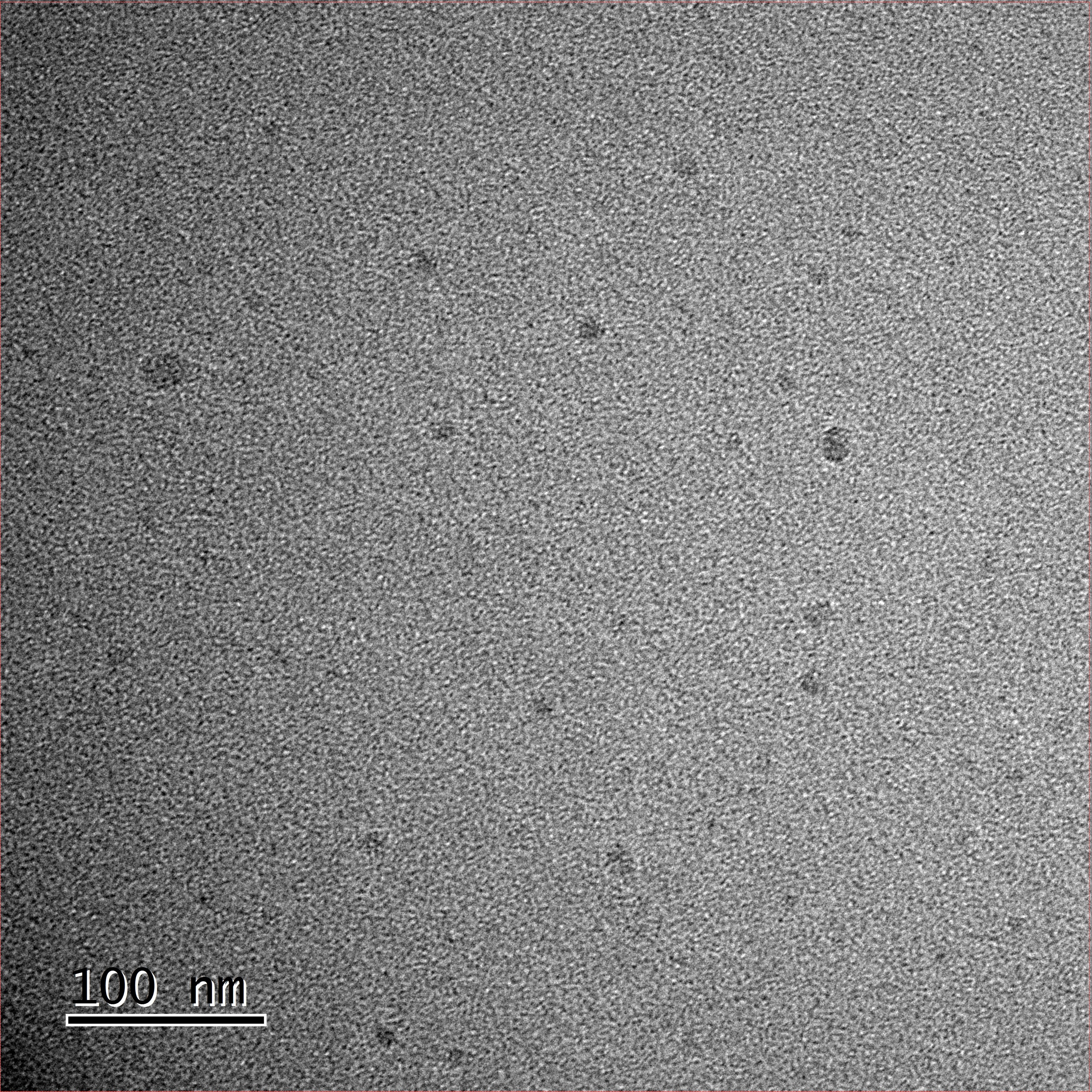

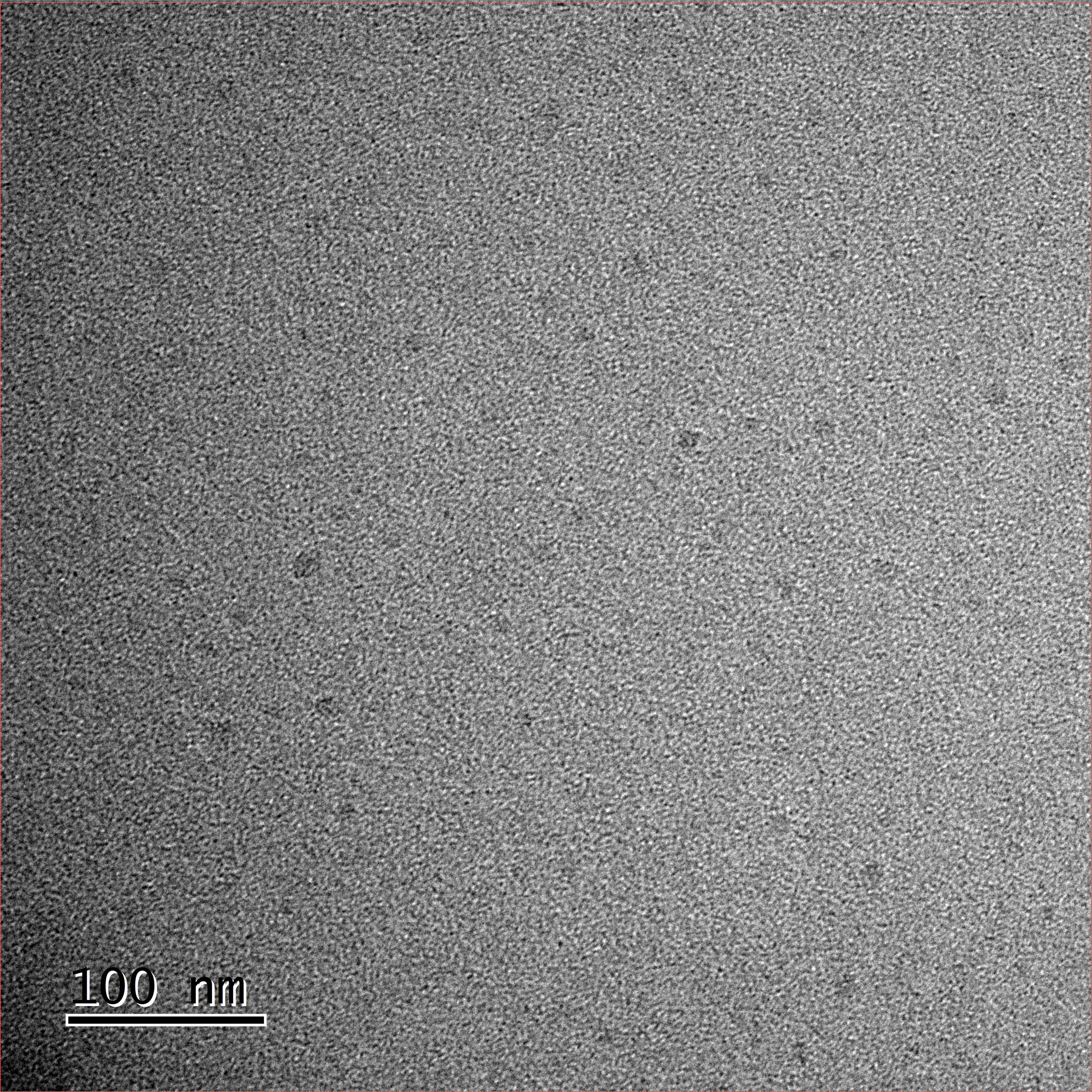

(a)

(b)

(c)

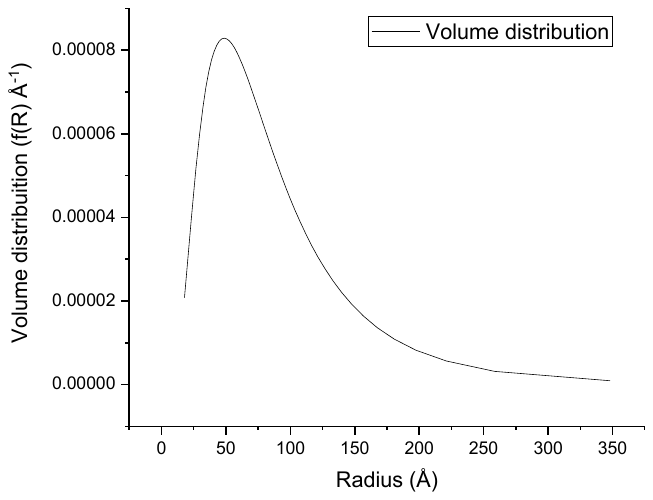

**Figure 7.** Volume distribution graph extracted from the raw SAXS data obtained for the MNPs, showing the mean radius as 9.07 nm.

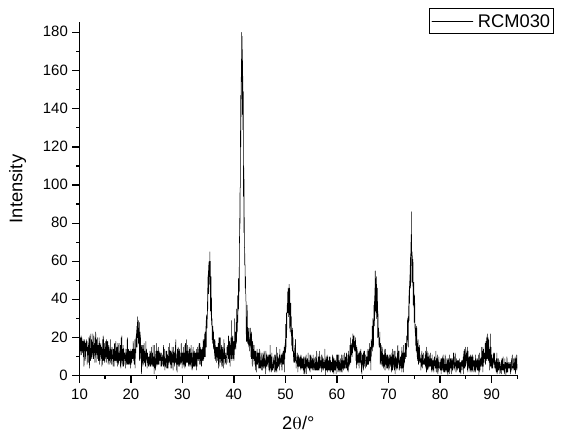

(111)

(220)

(311)

(400)

(222)

(422)

(511)

(440)

(531)

(620)

(533)

**Figure 8.** PXRD diagram of the MNPs, showing the Miller indices that match a literature-based spectrum of magnetite.

**Figure 9.** The interference assays of the IONPs and MNPs incubated with the cytotoxicity assay components, measured at the respective assays’ wavelengths. (a) IONPs incubated with NRU reagent for 3 hours and destain solution; (b) IONPs incubated with Resazurin reagent for 2 hours; (c) MNPs incubated with NRU for 3 hours and destain solution; (d) MNPs incubated with MTT reagent for 3 hours and DMSO; (e) MNPs incubated with Resazurin reagent for 2 and 3 hours.

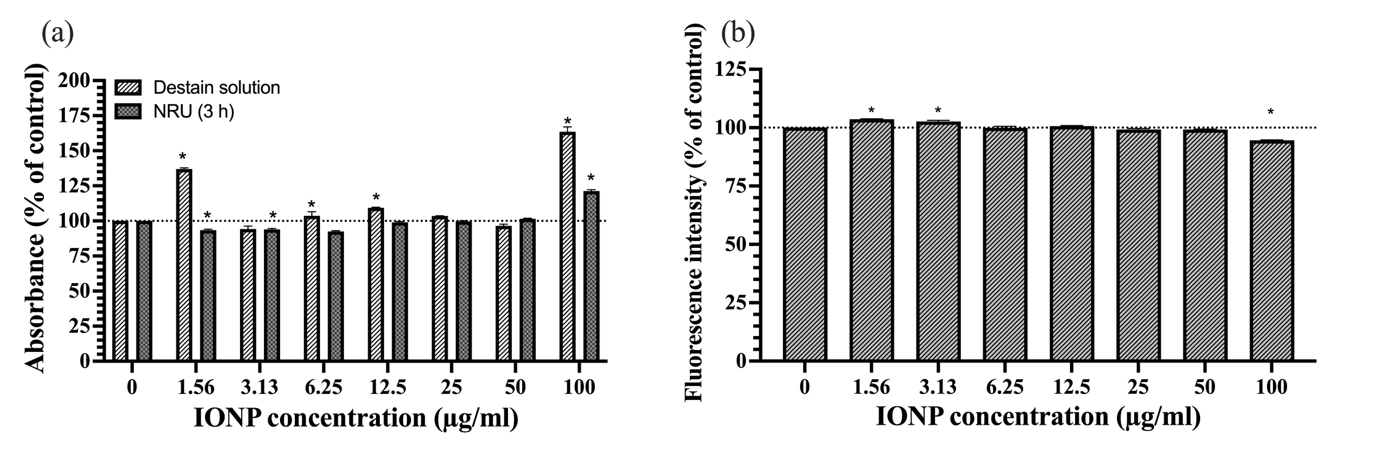

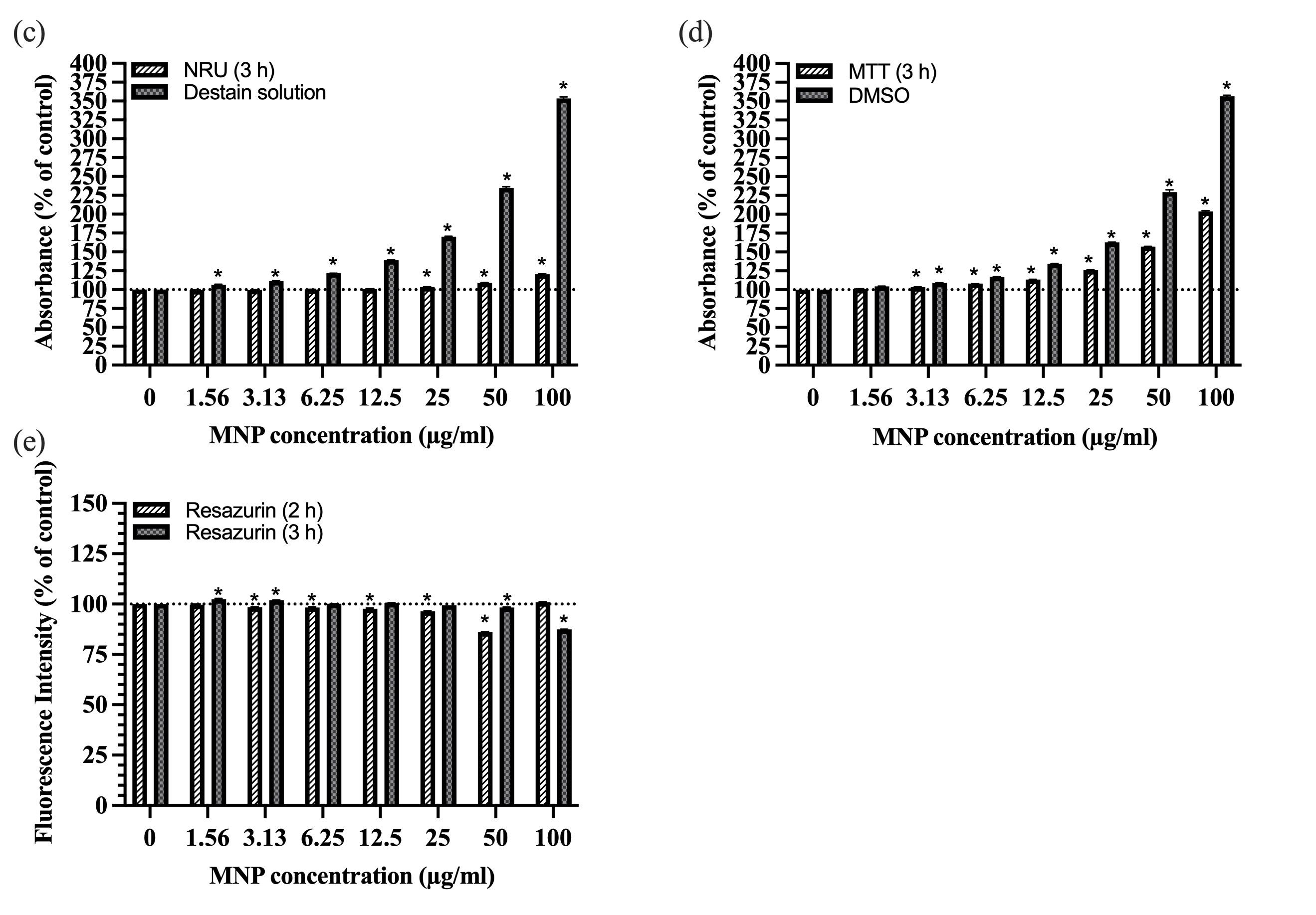

**Figure 10.** Remaining cell viability diagrams of the SK-N-SH and IMR-32 cell line exposures to IONPs for 24 hours. (a,b) Viability assessment of SK-N-SH and IMR-32 cells by NRU with fluorescence correction; (c,d) viability assessment of SK-N-SH and IMR-32 cells by Resazurin with absorbance correction. Bars represent standard error of the mean. Values were normalised to the positive and negative control, considering them as 0 and 100% respectively. *p < 0.05 represents a significant difference with respect to the negative control.

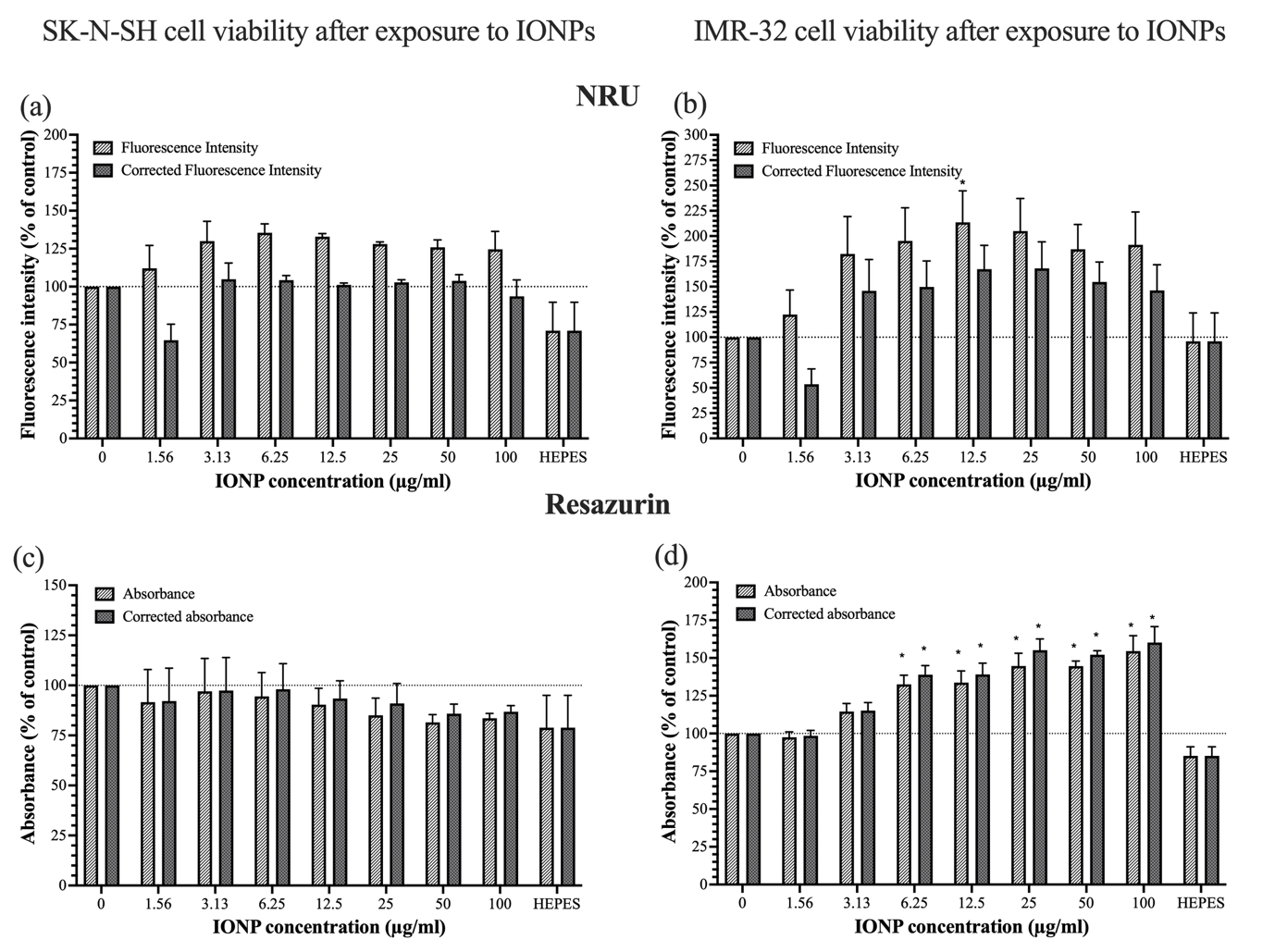

**Figure 11.** Remaining cell viability diagrams of the SK-N-SH and IMR-32 cell line exposures to MNPs for 4 hours. (a,b) Viability assessment of SK-N-SH and IMR-32 cells by NRU with pre- and post-transfer absorbance measurements; (c,d) viability assessment of SK-N-SH and IMR-32 cells by Resazurin with both reading modes. Bars represent standard error of the mean. Values were normalised to the positive and negative control, considering them as 0 and 100% respectively. *p < 0.05 represents a significant difference with respect to the negative control.

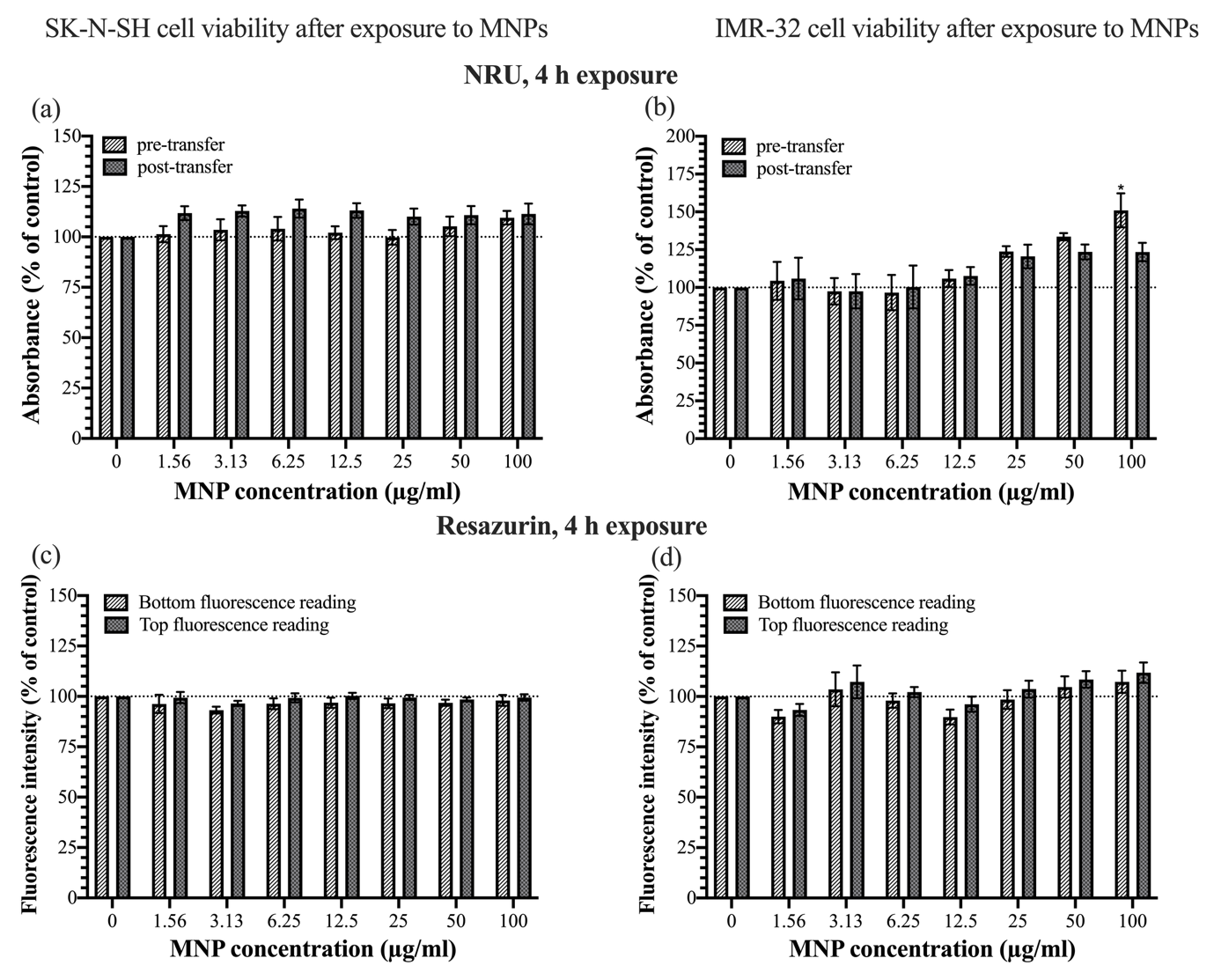
